# When the whole is less than the sum of its parts: maximum object category information and behavioral prediction in multiscale activation patterns

**DOI:** 10.1101/2021.07.14.452389

**Authors:** Hamid Karimi-Rouzbahani, Alexandra Woolgar

## Abstract

Neural codes are reflected in complex, temporally and spatially specific patterns of activation. One popular approach to decode neural codes in electroencephalography (EEG) is multivariate decoding. This approach examines the discriminability of activity patterns across experimental conditions to test if EEG contains information about those conditions. However, conventional decoding analyses ignore aspects of neural activity which are informative. Specifically, EEG data can be decomposed into a large number of mathematically distinct features (e.g., entropy, Fourier and Wavelet coefficients) which can reflect different aspects of neural activity. We previously compared 30 such features of EEG data, and found that visual category, and participant behavior, can be more accurately predicted using multiscale spatiotemporally sensitive Wavelet coefficients than mean amplitude (Karimi-Rouzbahani et al., 2021b). Here, we considered that even this larger set of features may only partially capture the underlying neural code, because the brain could use a combination of encoding protocols within a single trial which is not reflected in any one mathematical feature alone. To check, we combined those mathematical features using state-of-the-art supervised and unsupervised feature selection procedures (n = 17). Across 3 datasets, we compared decoding of visual object category between these 17 sets of combined features, and between combined and individual features. Object category could be robustly decoded using the combined features from all of the 17 algorithms. However, the combination of features, which were equalized in dimension to the individual features, were outperformed in most of the time points by the most informative individual feature (Wavelet coefficients). Moreover, the Wavelet coefficients also explained the behavioral performance more accurately than the combined features. These results suggest that a single but multiscale encoding protocol may capture the neural code better than any combination of features. Our findings put new constraints on the models of neural information encoding in EEG.

## Introduction

How is information about the world encoded by the human brain? Researchers have tried to answer this question using variety of brain imaging techniques across all sensory modalities. In vision, people have used invasive (Hung et al., 2005; ECoG; Majima et al., 2014; Watrous et al., 2015; Rupp et al., 2017; Lie et al., 2009; Miyakawa et al., 2018; Liu et al., 2009) and non-invasive (EEG and MEG; Contini et al., 2017; Carlson et al., 2013; Kaneshiro et al., 2016; Simanova et al., 2010; Cichy et al., 2014) brain imaging modalities to decode object category information from variety of features of the recorded neural activations. While majority of EEG and MEG decoding studies still rely on the within-trial “mean” of activity (average of activation level within the sliding analysis window) as the main source of information (Karimi-Rouzbahani et al., 2017b; Grootswagers et al., 2017), recent theoretical and experimental studies have shown evidence that temporal variabilities of neural activity (sample to sample changes in the level of activity) form an additional channel of information encoding (Orbán et al., 2016). For example, these temporal variabilities have provided information about the “complexity”, “uncertainty” and the “variance” of the visual stimulus, which correlated with the semantic category of the presented image (Garrett et al., 2020; Hermundstad et al., 2014; Orbán et al., 2016). Specifically, object categories which show a wider variability in their exemplars (e.g. houses) evoke more variable neural activation than categories which have lower variability (e.g. faces; Garrett et al., 2020). Accordingly, it is now clear that neural variabilities carry significant amounts of information about different aspects of sensory processing and may also play a major role in determining behavior (Waschke et al., 2021).

Despite the richness of information in neural variabilities, there is no consensus yet about how to quantify informative neural variabilities. Specifically, neural variabilities have been quantified using three classes of mathematical features: variance-, frequency- and information theory-based features, each detecting specific, but potentially overlapping aspects of the neural variabilities (Waschke et al., 2021). Accordingly, previous studies have decoded object category information from EEG using variance-based (Wong et al., 2006; Mazaheri and Jensen, 2008; Alimardani et al., 2018; Joshi et al., 2018), frequency-based (Watrous et al., 2015; Jadidi et al., 2016; Wang et al., 2018; Voloh et al., 2020; Taghizadeh-Sarabi et al., 2015; Wang et al., 2018) and information theory-based (Shourie et al., 2014; Richman et al., 2000; Ahmadi-Pajouh et al., 2018; Torabi et al., 2017) features. However, these previous studies remained silent about the temporal dynamics of category encoding as they performed the analyses (i.e. feature extraction and decoding) on the whole-trial data to maximize the decoding accuracy. On the other hand, time-resolved decoding analyses studied the temporal dynamics of category information encoding (Grootswagers et al., 2017; Kaneshiro et al., 2015; Karimi-Rouzbahani et al., 2018). However, few time-resolved studies have extracted any features other than the instantaneous activity at each time point, or the mean of activity across a short sliding window (e.g., by down-sampling the data), to incorporate the information contained in neural variabilities (Karimi-Rouzbahani et al., 2017a; Majima et al., 2014). Therefore, previous studies either did not focus on the temporal dynamics of information processing or did not include the contents of neural variabilities in time-resolved decoding.

Critically, as opposed to the Brain-Computer Interface (BCI) community, where the goal of feature extraction is to maximize the decoding accuracy, in cognitive neuroscience the goal is to find better neural correlates for the behavioral effect under study (Hebart and Baker, 2018; Williams et al., 2007; Jacobs et al., 2009; Woolgar et al., 2019; Karimi-Rouzbahani et al., 2021a; Karimi-Rouzbahani et al., 2021b). Specifically, a given feature is arguably only informative if it predicts behavior. Therefore, behavior is a key benchmark for evaluating the information content of any features including those which quantify neural variabilities. Interestingly, almost none of the above-mentioned decoding studies focused on evaluating the predictive power of their suggested informative features about behavior. Therefore, it remains unclear if the additional information they obtained from features of neural variabilities was task-relevant or epiphenomenal to the experimental conditions.

To overcome these issues, we proposed a new approach using medium-sized (50ms) sliding windows at each time step (5ms apart). The 50ms time window makes a compromise between concatenating the whole time window, which in theory allows any feature to be used at the expense of temporal resolution, and decoding in a time resolved fashion at each time point separately, which might lose temporal patterns of activity (Karimi-Rouzbahani et al., 2021b). Within each window, we quantify multiple different mathematical features of the continuous data. This allows us to be sensitive to any information carried in local temporal variability in the EEG response, while also maintaining reasonable temporal resolution in the analysis. In a recent study, we extracted a large set of such features and quantified the information contained in each using multivariate classification (Karimi-Rouzbahani et al., 2021b). We balanced the number of extracted values across features using Principal Component Analysis (PCA). Across three datasets, we found that that the incorporation of temporal patterns of activity in decoding, through the extraction of spatiotemporal “Wavelet coefficients” or even using the informative “original magnitude data (i.e. no feature extraction)”, provided higher decoding performance than the more conventional average of activity within each window (“mean”). Importantly, we also observed that for our Active dataset where participants categorized objects, the decoding results obtained from the same two features (i.e. Wavelet coefficients and original magnitude data) could predict/explain the participants’ reaction time in categorization significantly better than the “mean” of activity in each window (Wavelet outperformed original magnitude data). We further observed that more effective decoding of the neural codes, through the extraction of more informative features, corresponded to better prediction of behavioral performance. We concluded that the incorporation of temporal variabilities in decoding can provide additional category information and improved prediction of behavior compared to the conventional “mean” of activity.

One critical open question, however, is whether we should expect the brain to encode the information via each of these features individually, or whether it may instead use combinations of these features. In other words, while each of feature may potentially capture a specific and limited aspect of the generated neural codes, the brain may recruit multiple neural encoding protocols at the same time point or in succession within the same trial. Specifically, an encoding protocol might be active only for a limited time window or for specific aspects of the visual input (Victor, 2000; Wark et al., 2009; Gawne et al., 1996). For example, it has been shown in auditory cortex that two temporal encoding protocols (millisecond-order codes and phase coding) are simultaneously informative (Kayser et al., 2009). In visual stimulus processing, one study found that stimulus contrast was represented by latency coding at a temporal precision of ~10 ms, whereas stimulus orientation and its spatial frequency were encoded at a coarser temporal precision (30 ms and 100 ms, respectively; Victor, 2000). Another study showed that spike rates on 5-10 ms timescales carried complementary information to that in the phase of firing relative to low-frequency (1-8 Hz) LFPs about which epoch of naturalistic movie was being shown (Montemurro et al., 2008). Therefore, it might be the case that multiple neural encoding protocols contribute to the encoding of information. Alternatively, the brain may implement one flexible multiscale encoding protocol (e.g. multiplexing strategy which combines the codes at different time scales (Panzeri et al., 2010)), which allows different aspects of information to be represented within the same encoding protocol. This multiplexed encoding protocol has been suggested to provide several computational benefits including enhancing the coding capacity of the system (Kayser et al., 2009; Schaefer et al., 2006), reducing the ambiguity inherent to single-scale codes (Schaefer et al., 2006; Schroeder and Lakatos, 2009) and improving the robustness of neural representations to environmental noise (Kayser et al., 2009).

To see if the brain encodes information using several encoding protocols simultaneously, we created combinations from the large set of distinct mathematical features in our previous study (Karimi-Rouzbahani et al., 2021b). We asked whether their combination recovers more of the underlying neural code, leading to additional object category information and increased accuracy in predicting behavior, compared to the best performing individual feature from the previous study (i.e. Wavelet). Specifically, we used the same three datasets, extracted the same features from neural activity, selected the most informative features at each sliding time window and evaluated their information about object categories. We also evaluated how well each combined feature set explained behavioral recognition performance. Our prediction was that as targeted combinations of informative features provide more flexibility in detecting subtle differences, which might be ignored when using each individual feature, we should see both a higher decoding accuracy and predictive power for behavior compared to when using individual features. However, our results show that, the most informative individual feature (the Wavelet transform) outperformed all of the feature combinations (combined using 17 different feature selection algorithms). Similarly, Wavelet coefficients outperformed all combinations of features in predicting behavioral performance. Therefore, the brain seems to use a flexible multiscale encoding protocol (i.e. captured by Wavelet coefficients) rather than a combination of several encoding protocols simultaneously.

## Methods

As this study uses the same set of datasets and features used in our previous study, we only briefly explain the datasets and the features. The readers are referred to our previous paper (Karimi-Rouzbahani et al., 2021b) as well as the original papers (cited below) for more detailed explanation of the datasets and features. The datasets used in this study and the code are available online at https://osf.io/wbvpn/. The EEG and behavioral data are available in Matlab “.mat” format and the code in Matlab “.m” format.

All the open-source scripts used in this study were compared against other implementations of identical algorithms in simulations and used only if they produced identical results. All open-source implementation scripts of similar algorithms produced identical results in our simulations. To evaluate different implementations, we tested them using 1000 random (normally distributed with unit variance and zero mean) time series each including 1000 samples.

### Overview of Datasets

We selected three highly varied previously published EEG datasets (Table 1) for this study to be able to evaluate the generalizability of our results and conclusions. Specifically, the datasets differed in a wide range of aspects including the recording set-up (e.g. amplifier, number of electrodes, preprocessing steps, etc.), properties of the image-set (e.g. number of categories and exemplars within each category, colorfulness of images, etc.), paradigm and task (e.g. presentation length, order and the participants’ task). The EEG datasets were collected while the participants were presented with images of objects, animals, face, etc. Participants’ task in Dataset 1 was irrelevant to the identity of the presented objects; they reported if the color of fixation changed from the first stimulus to the second in pairs of stimuli. Participants’ task for Dataset 2 was to respond/withhold response to indicate if the presented object belonged to the category (e.g. animal) cued at the beginning of the block. Participants had no explicit active task except for keeping fixation on the center of the screen for Dataset 3. To obtain relatively high signal to noise ratios for the analyses, each unique stimulus was presented to the participants 3, 6 and 12 times in datasets 1, 2, 3, respectively. The three datasets previously succeeded in providing object category information using multivariate decoding methods. For more details about the datasets see the original manuscripts cited in Table 1.

**Table 1.**
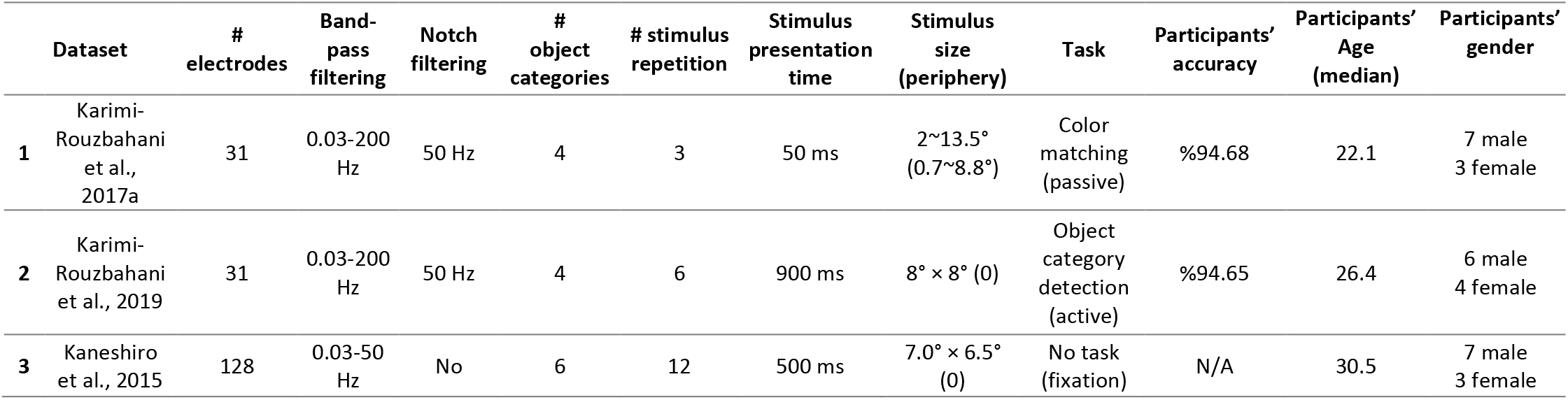
Details of the three datasets used in the study.

| Dataset | # electrodes | Band-pass filtering | Notch filtering | # object categories | # stimulus repetition | Stimulus presentation time | Stimulus size (periphery) | Task | Participants' accuracy | Participants' Age (median) | Participants' gender |
| --- | --- | --- | --- | --- | --- | --- | --- | --- | --- | --- | --- |
| **1** Karimi-Rouzbahani et al., 2017a | 31 | 0.03-200 Hz | 50 Hz | 4 | 3 | 50 ms | 2~13.5° (0.7~8.8°) | Color matching (passive) | %94.68 | 22.1 | 7 male<br>3 female |
| **2** Karimi-Rouzbahani et al., 2019 | 31 | 0.03-200 Hz | 50 Hz | 4 | 6 | 900 ms | 8° × 8° (0) | Object category detection (active) | %94.65 | 26.4 | 6 male<br>4 female |
| **3** Kaneshiro et al., 2015 | 128 | 0.03-50 Hz | No | 6 | 12 | 500 ms | 7.0° × 6.5° (0) | No task (fixation) | N/A | 30.5 | 7 male<br>3 female |

### Preprocessing

They were collected at a sampling rate of 1000 Hz. Each dataset consisted of data from 10 participants. Each object category in each dataset included 12 exemplars. For datasets 1 and 2, only the trials with correct responses were used in the analyses (dataset 3 had no task). To make the three datasets as consistent as possible, we pre-processed them differently from their original papers. Datasets 1 and 2 were band-pass-filtered in the range from 0.03 to 200 Hz. The band-pass filtering range of dataset 3 was 0.03 to 50 Hz, as we did not have access to the raw data to increase the upper bound to 200 Hz. We also performed notch-filtering on datasets 1 and two at 50 Hz. We used finite-impulse-response filters with 12 dB roll-off per octave for band-pass filtering of datasets 1 and 2 and when evaluating the sub-bands of the three datasets. We did not remove artifacts (e.g. eye-related and movement-related) from the signals as sporadic artifacts have minimal effect in multivariate decoding (Grootswagers et al., 2017). The filtering was applied on the data before they were epoched relative to the trial onset times. Data were epoched from 200 ms before to 1000 ms after the stimulus onset to cover most of the range of event-related neural activations. The average pre-stimulus (−200 to 0 ms relative to the stimulus onset) signal amplitude was removed from each trial of the data. For more information about each dataset see the references cited in Table 1.

### Features

We briefly explain the 26 mathematically distinct features used in this study below. Note that 4 of the features, which were event-related potentials, were excluded from this study as they could not be defined across time. For more details about their algorithms, their plausibility and possible neural underpinnings please see (Karimi-Rouzbahani et al., 2021b). Each feature was calculated for each EEG electrode and each participant separately.

Each of the following features was extracted from the 50 samples contained in 50 ms sliding time windows at a step size of 5 ms along each trial:

#### Mean, Variance, Skewness and Kurtosis

These are the standard 1st to 4th moments of EEG time series. To calculate these features, we simply calculated the mean, variance, skewness and variance of EEG signals ***over the samples within each sliding analysis window within each trial*** (50 samples). Please note that this differs from averaging over trials, which is sometimes used to increase signal to noise ratio (Hebart and Baker, 2018). “Mean” of activity is by far the most common feature of EEG signal used in time-resolved decoding (Grootswagers et al., 2017). Specifically, in time-resolved decoding, generally the samples within each sliding time window are averaged and used as the input for the classification algorithm. People sometimes perform down-sampling of EEG time series, which either performs simple averaging or retains the selected samples every few samples. Variance (Wong et al., 2006), Skewness (Mazaheri and Jensen, 2008) and Kurtosis (Alimardani et al., 2018; Pouryzdian and Erfaninan, 2010) have shown success in providing information about different conditions of visually evoked potentials.

#### Median

We also calculated signal’s median as it is less affected by spurious values compared to the signal mean providing less noisy representations of the neural processes.

While the moment features above provide valuable information about the content of EEG evoked potentials, many distinct time series could lead to similar moment features. In order to be sensitive to this potentially informative differences nonlinear features can be used which, roughly speaking, are sensitive to nonlinear and complex patterns in time series. Below we define the most common nonlinear features of EEG time series analysis, which we used in this study.

#### LZ complexity (LZ Cmplx)

We calculated the Lempel-Ziv complexity as an index of signal complexity. This measure counts the number of unique sub-sequences within the analysis window (50 time samples), after turning the time samples into a binary sequence. To generate the binary sequence, we used the signal median, within the same analysis window, as the threshold. Accordingly, the LZ complexity of a time series grows with the length of the signal and its irregularity over time. See (Lempel and Ziv, 1976) for more details. This measure has previously provided information about neural responses in primary visual cortices (Szczepański et al., 2003). We used the script by Quang Thai^1^ implemented based on “exhaustive complexity” which is considered to provide the lower limit of the complexity as explained by Lempel and Ziv (1976).

#### Higuchi and Katz fractal dimensions (Higuchi FD and Katz FD)

Fractal is an indexing technique which provides statistical information determining the complexity of how data are organized within time series. Accordingly, higher fractal values, suggest more complexity and vice versa. In this study, we calculated the complexity of the signals using two methods of Higuchi and Katz, as used previously for categorizing object categories (Namazi, 2018; Ahmadi-Pajouh et al., 2018; Torabi et al., 2017). We used the implementations by Jesús Monge Álvarez^2^ after verifying it against other implementations.

#### Hurst exponent (Hurst Exp)

This measure quantifies the long-term “memory” in a time series. Basically, it calculates the degree of dependence among consecutive samples of time series and functions similarly to the autocorrelation function (Racine, 2011; Torabi et al., 2017). Hurst values between 0.5 and 1 suggest consecutive appearance of high signal values on large time scales while values between 0 and 0.5 suggest frequent switching between high and low signal values. Values around 0.5 suggest no specific patterns among samples of a time series.

#### Sample and Approximate Entropy (Sample Ent and Apprx Ent)

Entropy measures the level of perturbation in time series. As the precise calculation of entropy needs large sample sizes and is also noise-sensitive, we calculated it using two of the most common approaches: sample entropy and approximate entropy. Sample entropy is not as sensitive to the sample size and simpler to implement compared to approximate entropy. Sample entropy, however, does not take into account self-similar patterns in the time series (Richman & Moorman, 2000). We used an open-source code^3^ for calculating approximate entropy.

#### Autocorrelation (Autocorr)

This index quantifies the self-similarity of a time series at specific time lags. Accordingly, if a time series has a repeating pattern at the rate of F hertz, an autocorrelation measure with a lag of 1/F will provide a value of 1. However, it would return −1 at the lag of 1/2F. It would provide values between −1 and 1 for other lags. More complex signals would provide values close to 0. A previous study has been able to decode neural information about motor imagery using the autocorrelation function from EEG signals (Wairagkar et al., 2016).

#### Hjorth Complexity and Mobility (Hjorth Cmp and Hjorth Mob)

These parameters measure the variation in the signals’ characteristics. The complexity measure calculates the variation in a signal’s dominant frequency, and the mobility measures the width of the signal’s power spectrum (how widely the frequencies are scattered in the power spectrum of the signal; (Joshi et al., 2018)).

#### Mean, Median and Average Frequency (Mean Freq, Med Freq and Avg Freq)

These measures calculate the central frequency of the signal in different ways. Mean frequency is the average of all frequency components available in a signal. Median frequency is the median normalized frequency of the power spectrum of the signal and the average frequency is the number of times the signal time series crosses zero. They have shown information about visual categories in previous studies (Iranmanesh and Rodriguez-Villegas, 2017; Joshi et al., 2018; Jadidi et al., 2016).

#### Spectral Edge Frequency (SEF 95%)

SEF indicates the high frequency below which x percent of the signal’s power spectrum exists. X was set to 95% in this study. Therefore, SEF reflects the upper-bound of frequency in the power spectrum.

#### Signal power, Power and Phase at Median Frequency (Signal Pw, Pw MdFrq and Phs MdFrq)

Power spectrum density (PSD) represents the intensity or the distribution of the signal power into its constituent frequency components. Signal power was used as a feature here as in previous studies (Rupp et al., 2017; Majima et al., 2014), where it showed associations between aspects of visual perception and power in certain frequency bands. Signal power is the frequency-domain representation of temporal neural variability (Waschke et al., 2021). We also extracted signal power and phase at median frequency which have previously shown to be informative about object categories (Jadidi et al., 2017; Rupp et al., 2017).

For the following features we had more than one value per trial and sliding time window. We extracted all these features but later down-sampled the values to ***one*** per trial using the (first) PCA procedure explained below (Figure 1) before using them in the feature combination procedure.

**Figure 1.**
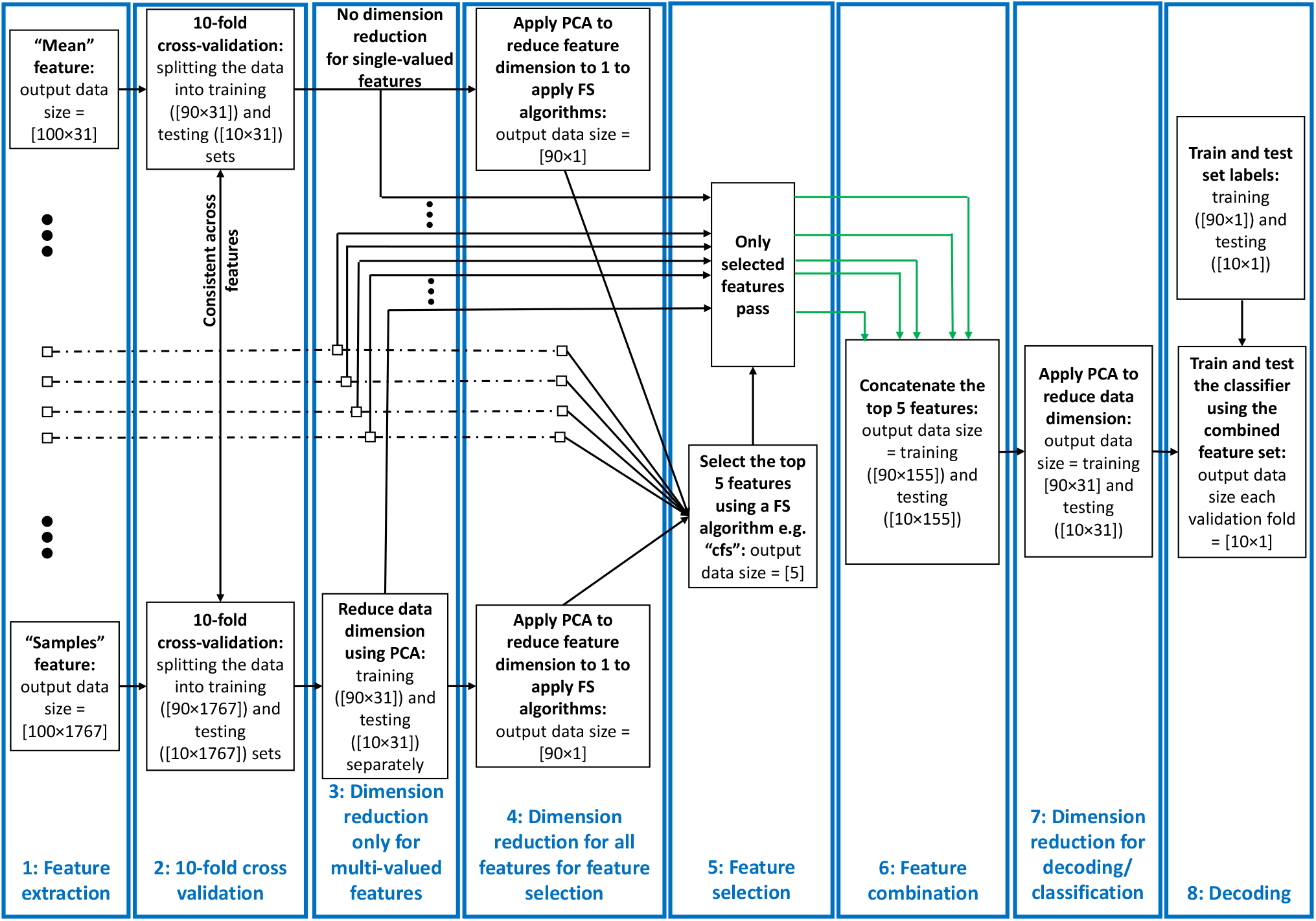
Decoding pipeline. From left to right: successive stages shown for a sample dataset comprising 100 trials of data from two categories recorded using a 31-electrode EEG amplifier. 1) Features are extracted from each trial and time window of the data. The features can be single- or multi-valued resulting in different number of values per trial and analysis time window. 2) We split the trials into training and testing sets and use the training sets in PCA and training the classifiers throughout the pipeline. 3) We used a PCA-based dimension reduction to reduce the number of values of only the multivalued features to one equalizing them with single-valued features. 4) We used a second PCA project all values of each feature to one dimension to be able to feed to the feature selection (FS) algorithms. 5) We selected the 5 most informative features using the FS algorithms. 6) We combined these features using concatenation of the selected features in their original size received from stage 4. 7) We reduced the dimension of the concatenated feature set to equalize it with the single-valued individual features from the previous study so that they could be compared. 8) We decoded/classified all pair-wise categories using the final dataset in each fold. This figure shows the procedure for a single cross-validation fold at one time point and was repeated for all the folds and time points. To avoid circularity, PCA was only ever applied on the training set and the parameters (mean and eigen vectors) used to derive the principal component of both the training and testing sets. The green arrows indicate example selected feature sets sent for combination.

#### Cross-correlation (Cros Cor)

This refers to the inter-electrode correlation of EEG time series. It simply quantifies the similarity of activations between pairs of EEG electrodes. Therefore, for each electrode, we had e-1 cross-correlation values with e referring to the number of electrodes. This measure has been shown to contain information about visual object categories before (Majima et al., 2014; Karimi-Rouzbahani et al., 2017a).

#### Wavelet Coefficients (Wavelet)

Considering the time- and frequency-dependent nature of ERPs, Wavelet transform seems to be a very reasonable choice as it provides a time-frequency representation of signal components. It determines the primary frequency components and their temporal position in time series. The transformation passes the signal time series through digital filters (Guo et al., 2009), each of which adjusted to extract a specific frequency (scale) at a specific time. This filtering procedure is repeated for several rounds (levels) filtering low- (approximations) and high-frequency (details) components of the signal to provide more fine-grained information about the constituent components of the signal. This can lead to coefficients which can potentially discriminate signals evoked by different conditions. Following up on a previous study (Taghizadeh-Sarabi et al., 2015), and to make the number of Wavelet features comparable in number to signal samples, we used detail coefficients at five levels D1,…,D5 as well as the approximate coefficients at level 5, A5. This led to 57 features in the 50 ms sliding time windows. We used the “Symlet2” basis function for our Wavelet transformations as implemented in Matlab. The multistage, variable-sized filtering procedure implemented in Wavelet coefficients, make them ideal for detecting multiscale patterns of neural activity, which has been suggested to be produced by the brain for information encoding (Panzeri et al., 2010).

#### Hilbert Amplitude and Phase (Hilb Amp and Hilb Phs)

This transformation is a mapping function that takes a function x(t) of a real variable, and using convolution with the function, 1/πt, produces another function of a real variable H(u)(t). This technique provides amplitude and phase information of the signal in the transformed space allowing us to tease them apart and evaluate their information content about visual categories (Wang et al., 2018).

#### Original magnitude data (Samples)

We also used the post-stimulus signal samples (i.e. 50 samples in each sliding analysis window) to decode object category information without any feature extraction. This allowed us to compare the information content of the extracted features with the original signal samples to see if the former provided any extra information. Note that, this is different from averaging/down-sampling of magnitude data within the analysis windows conventionally used in multivariate decoding (Karimi-Rouzbahani et al. 2017a).

### Feature selection algorithms

We set out to test whether neural information about object categories might be captured by combinations of the above features, better than by any one feature individually. For this, we combined the 26 extracted features using Feature Selection Library (FSLib, version 6.2.1; Roffo, 2016). Feature selection (FS), which refers to selecting a subset of features from a larger set, is generally used (for example, in machine learning) to reduce the dimensionality of the data by removing the less informative features from the dataset. FS algorithms can be categorized as supervised or unsupervised (Dash and Liu, 1997). The supervised methods receive, as input, the labels of trials for each condition (i.e. object categories here), and try to maximize the distance between conditions. We used 8 different supervised FS algorithms. The unsupervised methods, on the other hand, incorporate different criteria for FS such as selecting features that provide maximum distance (i.e. **unfol**) or minimum correlation (i.e. **cfs**). The FSLib implements 19 different feature selection algorithms. As it is not yet known how the brain might recruit different encoding protocols or a potential combination of them, we used all the FS algorithms available by the FSLib to combine the features in this study, except two (rfe-SVM and L0) which we were not able to implement. We set the number of selected features to 5, which was chosen to balance between including too many features, which could obscure interpretability, and including too few, which risks missing informative but lower-ranked features. Below we briefly explain the 8 supervised and 9 unsupervised feature selection algorithms. Readers are referred to the original papers for more detail about each feature selection method as reviewed (Roffo, 2016).

Among supervised algorithms, ***Relief*** is a randomized and iterative algorithm that evaluates the quality of the features based on how well their values discriminate data samples from opposing conditions. This algorithm can be sensitive when used on small data samples. ***Fisher*** evaluates the information of features as the ratio of inter-class to intra-class distances. ***Mutual Information*** (mutinffs) measures the association between the data samples (observations) within each feature and their class labels. ***Max-Relevance, Min-Redundancy*** (mrmr) method, which is an extension of the mutual information method, is designed to follow two basic rules when selecting the features: to select the features which are mutually far away from each other while still having “high” correlation to the classification labels. As opposed to the above methods, which rank and select the features according to their specific criteria, the ***Infinite latent*** (ILFS) method, selects the most informative features based on the importance of their neighboring features in a graph-based algorithm. It is a supervised probabilistic approach that models the features “relevancy” in a generative process and derives the graph of features which allows the evaluation of each feature based on its neighbors. Similarly, the method of ***Eigenvector Centrality*** (ECFS), generates a graph of features with features as nodes and evaluates the importance of each node through an indicator of centrality i.e. eigen vector centrality. The ranking of central nodes determines the most informative features. ***LASSO*** algorithm works based on error minimization in predicting the class labels using the features as regression variables. The algorithm penalizes the coefficients of the regression variables while setting the less relevant to zero to follow the minimal sum constraint. The selected features are those which have non-zero coefficients in this process. ***Concave Minimization*** (fsv) uses a linear programming technique to inject the feature selection process into the training of a support vector machine (SVM).

Among unsupervised FS algorithms, ***Infinite FS*** (InfFS), is similar to the graph-based supervised methods in which each feature is a node in a graph. Here, however, a path on a graph is a subset of features and the importance of each feature is measured by evaluating all possible paths on the graph as feature subsets in a cross-validation procedure. ***Laplacian Score*** (laplacian), evaluates the information content of each feature by its ability of locality preserving. To model the local geometry of the features space, this method generates a graph based on nearest neighbor and selects the features which respect this graph structure. ***Dependence Guided*** (dgufs) method evaluates the relationship between the original data, cluster labels and selected features. This algorithm tries to achieve two goals: to increase the dependence on the original data, and to maximize the dependence of the selected features on cluster labels. ***Adaptive Structure Learning*** (fsasl), which learns the structure of the data and FS at the same time is based on linear regression. ***Ordinal Locality*** (ufsol) is a clustering-based method which achieves distance-based clustering by preserving the relative neighborhood proximities. ***Multi-Cluster*** (mcfs) method is based on manifold learning and L1-regularized models for subset selection. This method selects the features such that the multi-cluster structure of the data can be best preserved. As opposed to most of the unsupervised methods which try to select the features which preserve the structure of the data, e.g. manifold learning, ***L2,1-norm Regularized*** (UDFS) method assumes that the class label of data can be predicted using a linear classifier and incorporates discriminative analysis and L2,1-norm minimization into a joint framework for feature selection. ***Local Learning-Based*** (llcfs) method is designed to work with high-dimensional manifold data. This method associates weights to features which are incorporated into the regularization procedure to evaluate their relevance for the clustering. The weights are optimized iteratively during clustering which leads to the selection of the most informative features in an unsupervised fashion. ***Correlation-Based*** (cfs) method simply ranks the features based on how uncorrelated they are to the other features in the feature set. Therefore, the selected features are those which are most distinct from others.

### Decoding pipeline

The pipeline used in this study for feature extraction, dimensionality reduction, feature selection, feature combination and decoding had 8 stages and is summarized in Figure 1. Below we explain each stage of the pipeline for a simple sample dataset with 100 trials collected using a 31-electrode EEG setup. Our actual datasets, however, had varied number of trials and electrodes as explained above.

#### 1: Feature extraction

We extracted the set of 26 above-mentioned features from the dataset. This included features which provided one value for each sliding time window per trial (single-valued) and more than one value (multi-valued). For the sample dataset, this resulted in data matrices with 100 rows (trials) and 31 columns (electrodes) for the single-valued datasets and 31× *e* columns for multi-valued features, where *e* refers to the number of values extracted for each trial and time window.

#### 2: Cross validation

After extracting the features, we split the data into 10 folds, used 9 folds for dimension reductions and training the classifiers and the left-out fold for testing the classifiers. Therefore, we used a 10-fold cross-validation procedure in which we trained the classifier on 90% of the data and tested it on the left-out 10% of the data, repeating the procedure 10 times until all trials from the pair of categories participate once in the training and once in the testing of the classifiers. The same trials were chosen for all features in each cross-validation fold.

#### 3: Dimensionality reduction 1: only for multi-valued features

The multi-valued features explained above resulted in more than a single feature value per trial per sliding time window (e.g. cross-correlation, wavelet, Hilbert amplitude and phase and signal samples). This could lead to the domination of the multi-valued over single-valued features in feature selection and combination. To avoid that, we used principle component analysis (PCA) to reduce the number of values in the multi-valued features to one per electrode per time window, which was the number of values for all single-valued features. Specifically, the data matrix before dimension reduction, had a dimension of *n* rows by *e* × *f* columns where *n, e* and *f* were the number of trials in the dataset (consisting of all trials from all categories), the number of electrodes and the number of values from obtained from a given feature (concatenated in columns), respectively. *Therefore, the columns of multi-valued features included both the **spatial (electrodes)** and **temporal (elements of each feature)** patterns of activity from which the information was obtained. This is different from single-valued features where the columns of their data matrix only included **spatial** patterns of activity*. As *f* = 1 for the single-valued features, for the multi-valued features, we only retained the *e* most informative columns that corresponded to the *e* eigen values with highest variance and removed the other columns using PCA. Therefore, we reduced the dimension of the data matrix to *n* × *e* which was the same for single- and multi-valued features and used the resulting data matrix for decoding. This means that, for the multi-valued features, in every analysis window, we only retained the most informative value of the extracted feature elements and electrode (i.e. the one with the most variance in PCA). Accordingly, multi-valued features had the advantage over single-valued features as the former utilized both the ***spatial*** and ***temporal*** patterns of activity in each sliding time window, while the latter only had access to the ***spatial*** patterns.

#### 4: Dimensionality reduction 2: for feature selection

For feature selection, each feature should have a dimension of 1 to go into the FS algorithm. However, our features had as many dimensions as the number of electrodes (i.e. *e*). Therefore, we further reduced the dimension of each feature from *e* to 1 to be able to feed them to the FS algorithms, compare them and select the most informative features. This allowed us to know the general amount of information that each feature rather than each of its elements/dimensions (e.g. electrodes in single-valued features) had about object categories. Please note that, however, after finding the most informative features, we used the selected features in their original size which was *e* (output of step 3 goes to stage 6).

#### 5: Feature selection

Feature selection was done using 17 distinct algorithms (above) to find the 5 most informative features in every sliding time window. This stage only provided indices of the selected features for combination in the next stage. To avoid any circularity (Pulini et al., 2019), we applied the FS algorithms only on the training data (folds) and used the selected features in both training and testing in each cross-validation run. Please note that feature selection was performed in every analysis window across the trial. In other words, different sets of 5 features could be selected for each individual analysis window. This allowed multiple features to contribute at each time point (multiple codes to be in use at the same time) and for different features to be selected at different time points (different codes used at different points in the trial).

#### 6: Feature combination

We only concatenated the 5 selected features into a new data matrix. At this stage, we received 5 feature data matrices which had a dimension of *n* × *e* with *n* referring to the number of trials and *e* referring to the number of values per trial, which were 100 × 31 for the sample dataset explained in Figure 1. The combination procedure led to a concatenated data matrix of 100 × 155 (*n* × 5*e*).

#### 7: Dimensionality reduction 3: equalizing the dimensions of combined and individual feature spaces

We used another round of PCA to simultaneously combine and reduce the dimensionality of each data matrix (feature space) to equalize it with the feature space of the individual features. This made the combined and individual features directly comparable, so that we could test whether a combination of the most informative features could provide additional category-related information, over and above the information decodable from individual features. Had we not controlled for the dimension of the data matrix, superior decoding for the combined features could arise trivially (due to having more predictors). Note that, whereas we knew the features which were selected on stage 5, as a result of this PCA transformation, we did not know which features contributed to the final decoding result. Therefore, in the worst case scenario, the final feature set might have only contained one of the 5 selected features. However, this seems unlikely to be the case as generally all inputs contribute to the distributions of the data in the PCA space. To avoid circularity (Pulini et al., 2019), we again applied the PCA algorithms on the training data (folds) only and used the training PCA parameters (i.e. eigen values and means) for both training and testing (fold) sets for dimension reduction, carrying this out in each cross-validation run separately.

#### 8: Multivariate decoding

Finally we used time-resolved multivariate decoding to test for information about object categories in the features and combinations of features. We used linear discriminant analysis (LDA) classifiers to measure the information content across all possible pairs of conditions (i.e. object categories) in each dataset. We repeated the decoding across all possible pairs of categories within each dataset, which were 6, 6 and 15 pairs for datasets 1, 2 and 3, which consisted of 4, 4 and 6 object categories, respectively. Finally, we averaged the results across all combinations and reported them as the average decoding for each participant. We extracted the features from 50 ms sliding time windows in steps of 5 ms across the time course of the trial (−200 to 1000 ms relative to the stimulus onset time). Therefore, the decoding results at each time point reflect the data for the 50 ms window around the time point, from −25 to +24 ms relative to the time point.

### Decoding-behavior correlation

We evaluated the correlation between neural representations of object categories and the reaction time of participants in discriminating them. To that end, we generated a 10-dimensional vector of neural decoding accuracies (averaged over all pairwise category decoding accuracies obtained from each participant) at every time point and a 10-dimensional vector which contained the behavioral reaction times (averaged over all categories obtained from each participant) for the same group of 10 participants. Then we correlated the two vectors at each time point using Spearman’s rank-order correlation (Ritchie et al., 2015; Cichy et al., 2014). This resulted in a single correlation value for each time point for the group of 10 participants.

### Parameters of decoding curves

To quantitatively evaluate the patterns of decoding curves and decoding-behavior correlations, we extracted four distinct parameters from the decoding curves and one parameter from the correlation to behavior curves. All parameters were calculated in the post-stimulus time span. The “average correlation to behavior” was calculated by averaging the level of across-subject correlation to behavior. The parameters of “average decoding” and “maximum decoding” were calculated for each participant simply by calculating the average and maximum of the decoding curves. The “time of maximum decoding” and “time of first above-chance decoding” were also calculated for each participant relative to the time of the stimulus onset.

### Statistical analyses

#### Bayes factor analysis

First we asked whether we could decode object category from the combined features returned by each of the 17 FS methods. To determine the evidence for the null and the alternative hypotheses, we used Bayes analyses as implemented by Bart Krekelberg^4^ based on Rouder et al. (2012). We used standard rules of thumb for interpreting levels of evidence (Lee and Wagenmakers, 2014; Dienes, 2014): Bayes factors of >10 and <1/10 were interpreted as strong evidence for the alternative and null hypotheses, respectively, and >3 and <1/3 were interpreted as moderate evidence for the alternative and null hypotheses, respectively. We considered the Bayes factors which fell between 3 and 1/3 as suggesting insufficient evidence either way.

To evaluate the evidence for the null and alternative hypotheses of at-chance and above-chance decoding, respectively, we compared the decoding accuracies obtained from all participants in the post-stimulus onset time against the decoding accuracies obtained from the same participants averaged in the pre-stimulus onset time (−200 to 0 ms). We also asked whether there was a difference between the decoding values obtained from all possible pairs of FS methods. Accordingly, we performed the Bayes factor unpaired *t-test* and calculated the Bayes factor as the probability of the data under alternative (i.e. difference; H1) relative to the null (i.e. no difference; H0) hypothesis between all possible pairs of FS methods for each dataset separately. The same procedure was used to evaluate evidence for difference (i.e. alternative hypothesis) or no difference (i.e. null hypothesis) in the maximum and average decoding accuracies, the time of maximum and above-chance decoding accuracies across FS methods for each dataset separately. To evaluate the evidence for the null or alternative hypotheses of lack of or the existence of difference between the decoding accuracies obtained from FS algorithm and the Wavelet feature, we calculated the Bayes factor between the distribution of the two distributions of decoding accuracies on every time point and for dataset separately.

The priors for all Bayes factor analyses were determined based on Jeffrey-Zellner-Siow priors (Jeffreys, 1961; Zellner and Siow, 1980) which are from the Cauchy distribution based on the effect size that is initially calculated in the algorithm using t-test (Rouder et al., 2012). The priors are data-driven and have been shown to be invariant with respect to linear transformations of measurement units (Rouder et al., 2012), which reduces the chance of being biased towards the null or alternative hypotheses. No correction for multiple comparisons have been performed when using Bayes factors as they are much more conservative than frequentist analysis in providing false claims with confidence (Gelman and Tuerlinckx, 2000; Gelman et al., 2012). The reason for the less susceptibility of Bayesian analysis compared to classical statistics, is the use of priors, which if chosen properly (here using the data-driven approach developed by Rouder et al. (2012)), significantly reduce the chance of making type I (false positive) errors.

#### Random permutation testing

To evaluate the significance of correlations between decoding accuracies and behavioral reaction times, we calculated the percentage of the actual correlations that were higher (if positive) or lower (if negative) than a set of 1000 randomly generated correlations. These random correlations were obtained by randomizing the order of participants’ data in the behavioral reaction time vector (null distribution) on every time point, for each feature separately. The correlation was considered significant if surpassed 95% of the randomly generated correlations in the null distribution in either positive or negative directions (p < 0.05) and the p-values were corrected for multiple comparisons across time using Matlab mafdr function, where the algorithm fixes the rejection region and then estimates its corresponding error rate resulting in increased accuracy and power (Storey, 2002).

## Results

### Do different ways of combining individual features affect the level and temporal dynamics of information decoding?

As an initial step, we evaluated the level of information which can be obtained from the combination of features, each potentially capturing different aspects of the neural codes. To be as confident as possible, we used a large set of 17 distinct supervised and unsupervised feature selection (FS) methods to select and combine the top 5 most informative features at every time point in the time-resolved decoding procedure. The information content of features were determined based on either how much they could contribute to discriminating the target object categories (supervised) or some predefined criteria which could implicitly suggest more separation between object categories (unsupervised). We split the FS algorithms into three arbitrary groups for the sake of clearer presentation of the results (Figure 2).

**Figure 2.**
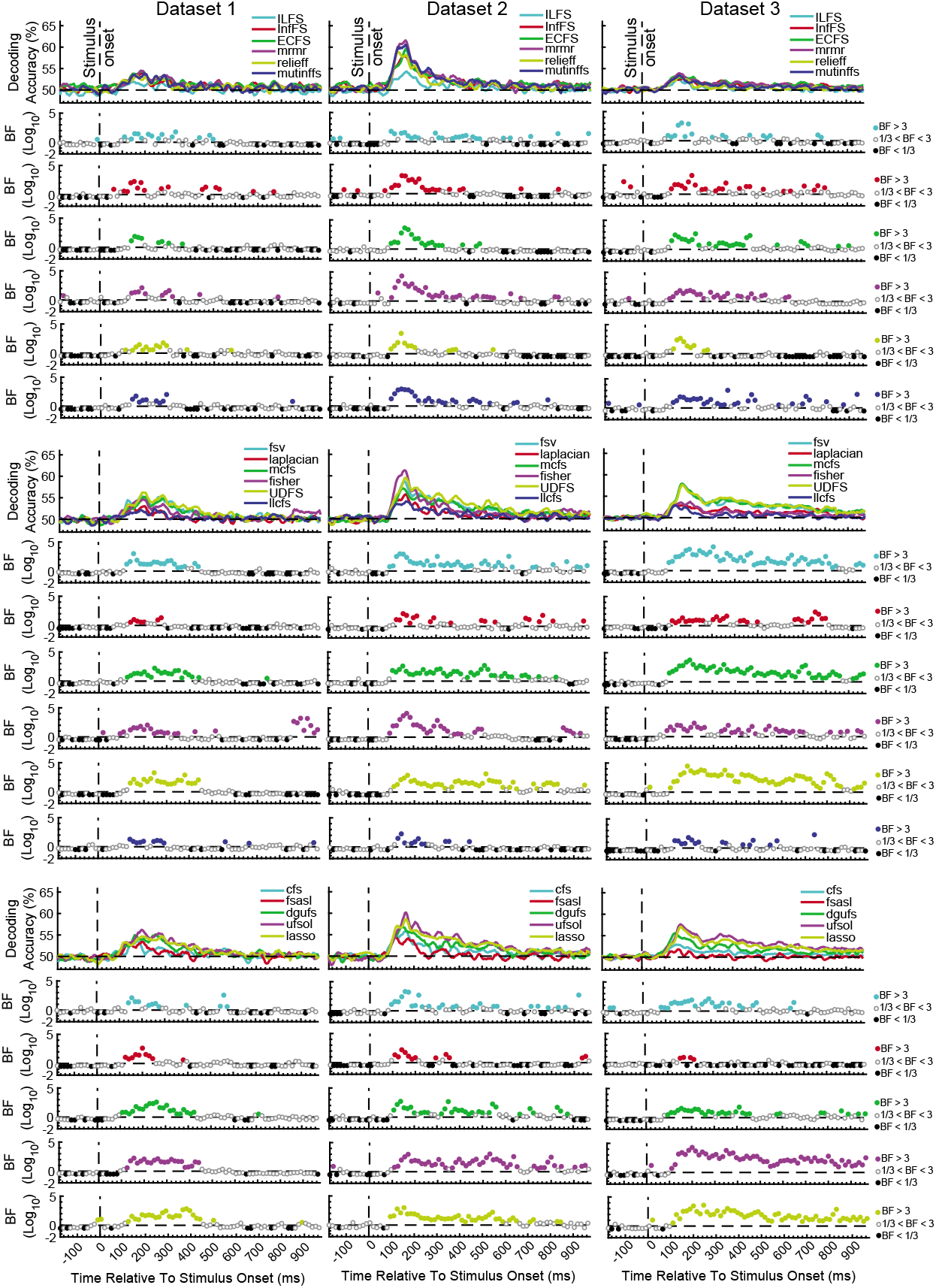
Time-resolved decoding of object categories from the three datasets using the 17 feature selection (FS) methods. We split the FS algorithms into three arbitrary groups (rows) for each dataset for the sake of clearer presentation. Each column shows the results for one dataset. The top section in each of the nine panels shows the decoding accuracies across time and the bottom panels show the Bayes factor evidence for decoding to be different (H1) or not different (H1) from chance-level. The horizontal dashed lines refer to chance-level decoding, the vertical dashed lines indicates time of stimulus onset. Non-black colored filled circles in the Bayes Factors show moderate (BF>3) or strong (BF>10) evidence for difference from chance-level decoding, black filled circles show moderate (BF>3) or strong (BF>10) evidence for no difference from chance-level decoding and empty circles indicate insufficient evidence (1/3<BF<3) for either hypotheses.

All FS algorithms for the three datasets showed strong (BF>10) evidence for difference from chance-level decoding at some time points/windows after the stimulus onset (Figure 2). This means that, any of the FS algorithms could combine the features in a way that they could decode object category information from brain signals. As expected from the difference in their mathematical formulations, however, no pairs of FS algorithms provided identical patterns of decoding in any of the three datasets. Consistently across the three datasets there was moderate (3<BF<10) or strong (BF>10) evidence for continuous above-chance decoding from around 80 ms post stimulus onset for all FS algorithms. While the decoding showed evidence for above-chance accuracy (BF>3) up until 550 ms (dataset 2) or even later than 800 ms (dataset 3) for the best FS algorithms such as UDFS, lasso and ufsol, all curves converged back to the chance-level earlier than 500 ms for dataset 1. This difference may reflect the longer stimulus presentation time for datasets 2 and 3 vs. dataset 1, which may have provided stronger sensory input for neural processing of category information, as we saw previously when evaluating individual features alone (Karimi-Rouzbahani et al., 2021b).

In order to quantitatively compare the decoding curves for the different FS algorithms, we extracted four different amplitude and timing parameters from their decoding curves as in previous studies (Isik et al., 2014): maximum and average decoding accuracies (in the post-stimulus time window), time of maximum decoding, and time of first above-chance decoding relative to stimulus onset (Supplementary Figure 1). Results showed that ILFS, relief and llcfs were the worst performing FS algorithms with the lowest maximum and average decoding accuracy (Supplementary Figure 1A and B; red boxes). UDFS, lasso and ufsol were the best performing FS algorithms leading to the highest maximum and average decoding accuracies (Supplementary Figure 1A and B; black boxes). Dataset 2 tended to yield higher decoding accuracies compared to the other datasets, which might be attributed to the longer presentation time of the stimuli and the active task of the participants, (Karimi-Rouzbahani et al., 2021a; Karimi-Rouzbahani et al., 2021c; Roth et al., 2020). UDFS, ufsol and relief were among the earliest FS algorithms to reach their first above-chance and maximum decoding accuracies (Supplementary Figure 1C and D). However, there was not a consistent pattern of temporal precedence for any FS algorithms across the datasets.

### Which individual features are selected by the most successful algorithms?

The difference in the decoding patterns for different FS algorithms suggest that they used different sets of features in decoding. To see what features were selected by different FS algorithms, and whether the informative individual features were selected, we calculated the merit of each of the individual features in each FS algorithm across the time course of the trial (Supplementary Figure 2). Here, merit refers to the frequency of a feature being selected by the FS algorithm for decoding. We calculated the merit as the ratio of the number of times the feature was among the top selected 5 features to the number of times the decoding was performed on every time point (i.e. all possible combination of category pairs).

Visual inspection of the results suggests that each FS algorithm seemed to rely on consistent sets of features across the three datasets, which are generally different between FS algorithms. This reflects that different FS algorithms have different levels of sensitivity and distinct selection criteria. Results also showed that the merit of different features varied across the time course of trials based on their information content about object categories relative to other features (Supplementary Figure 2). Therefore, the recruitment of features varied across the time course of the trial: while some features were only temporarily selected (e.g. Average and Mean frequency in the laplacian method from ~200 to 600 post-stimulus onset), there were features which were constantly used for decoding even before the stimulus onset (e.g. Cros Cor in the fsasl method), although they did not lead to any information decoding in the pre-stimulus time span (Figure 2). This might again be explained by the different levels of sensitivity and distinct selection criteria implemented by different FS algorithms. Importantly, the FS algorithms that provided the highest level of decoding (i.e. ufsol, lasso and UDFS) showed the highest merits for the features of Mean, Median, Samples and Wavelet which were among the most informative features when evaluated individually across the three datasets (Karimi-Rouzbahani et al., 2021b). On the other hand, the FS algorithms that performed most poorly (ILFS, releiff and llcfs) either used scattered sets of features (ILFS) or did not use the informative features of Mean, Median, Samples and Wavelet (llcfs and relief). Therefore, the FS algorithms that used the informative individual features outperformed other FS algorithms which did not.

### Are the neural codes better captured by a combinatorial encoding protocol or by a flexible multiscale encoding protocol?

The main question of this study was to see whether the flexibility obtained by the combination of features provides any additional information about object categories compared to the best-performing individual features by detecting the neural codes more completely. In other words, we wanted to test the hypothesis that the brain uses a combination of different neural encoding protocols simultaneously as opposed to using a flexible multiscale encoding protocol (such as reflected in the Wavelet transform). To test this hypothesis, we directly compared the decoding accuracy obtained from the top performing individual feature from the original study (Wavelet; Karimi-Rouzbahani et al., 2021b), which is able to detect multiscale spatiotemporal patterns of information, with the decoding accuracy obtained from the top performing FS algorithm, which used a set of combined features (ufsol; Figure 3). Results showed consistent patterns across the three datasets with the Wavelet feature outperforming the decoding accuracies obtained by the ufsol FS algorithm across most time points. Maximum continuous evidence for difference (BF>10) occurred between 80 to 320 ms, 75 to 180 ms and 85 to 325 ms for datasets 1-3, respectively. Therefore, it seems that, at least for object categories, the coding scheme in the brain is best captured by a flexible multiscale encoding protocol (implemented here by the Wavelet coefficients), rather than a combination of distinct encoding protocols (captured here by different features).

**Figure 3.**
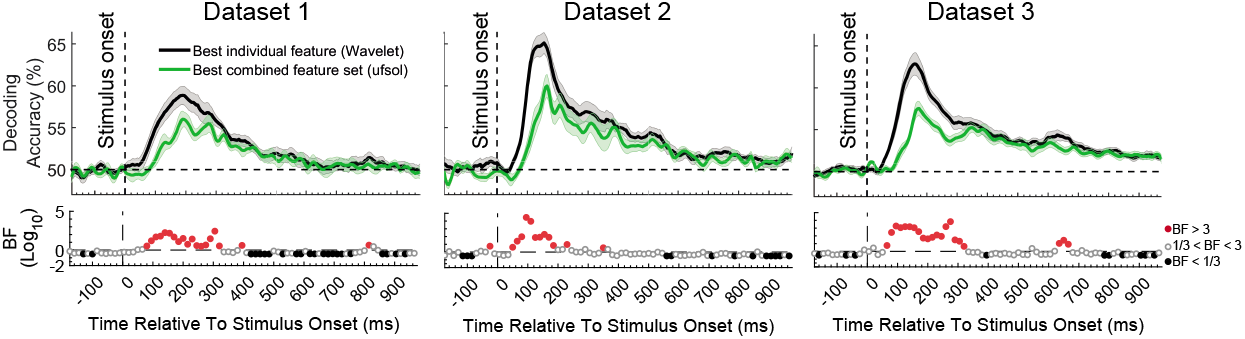
Comparison of decoding accuracies between the most informative individual feature (Wavelet; from Karimi-Rouzbahani et al., 2021b) and combined feature set (obtained using ufsol algorithm) from the three datasets and Bayesian evidence for a difference between them. Each column shows the results for one dataset. Thick lines show the average decoding accuracy across participants (error bars show Standard Error across participants). Top section in each panel shows the decoding accuracies across time and the bottom section shows the Bayes factor evidence for the difference between the decoding curves. The horizontal dashed lines on the top panels refer to chance-level decoding. Red filled circles in the Bayes Factors show moderate (BF>3) or strong (BF>10) evidence for difference between decoding curves, black filled circles show moderate (BF>3) or strong (BF>10) evidence for no difference and empty circles indicate insufficient evidence (1/3<BF<3) for either hypotheses.

### Can a combinatorial encoding protocol predict behavioral accuracy better than a flexible multiscale encoding protocol?

Our final hypothesis was that a combinatorial encoding protocol might predict the behavioral performance more accurately than a flexible multiscale encoding protocol as the former can potentially detect more distinctly encoded neural codes from brain activation. We could test this hypothesis only for Dataset 2 where the task was active and we had the participants’ reaction times (i.e. time to categorize objects) to work with. We calculated the (Spearman’s rank) correlation between the decoding accuracies and the behavioral reaction time across participants, to see whether, at each time point, participants with higher decoding values were those with the fastest reaction times. We expected to observe negative correlations between the decoding accuracies and the participants’ reaction times in the post-stimulus span (Ritchie et al., 2015). Note that since correlation normalizes the absolute level of the input variables, the higher level of decoding for the individual (Wavelet) feature vs. the combined features (ufsol; Figure 3) does not necessarily predict a higher correlation for the individual feature of Wavelet.

Results showed significant negative correlations appearing after the stimulus onset for most FS algorithms (except dgufs) especially the laplacian algorithm which showed the most negative peak (Figure 4A). This confirms that the distances between object categories in neural representations have inverse relationship to behavioral reaction times (Ritchie et al., 2015). We previously observed that the individual features which provided the highest decoding accuracies could also predict the behavior most accurately (Karimi-Rouzbahani et al., 2021b). Therefore, we asked if the FS algorithms which provided the highest levels of decoding could also predict the behavior more accurately than the less informative algorithms. The rationale behind this hypothesis was that, more effective decoding of neural codes, as measured by higher “average decoding” and “maximum decoding” accuracies (Figure 2), should facilitate the prediction of behavior by detecting subtle but overlooked behavior-related neural codes. To test this hypothesis, we evaluated the correlation between the parameters of “maximum decoding” and “average decoding” accuracies (extracted from the decoding curve of each feature in Figure 4A) and the “average correlation to behavior” (calculated simply by averaging the correlation to behavior in the post-stimulus time span for each FS algorithm in Figure 4A). We also calculated the correlation between the “time of maximum decoding” and “time of first above-chance decoding” as control variables, which we did not expect to correlate with behavior (as in Karimi-Rouzbahani et al., 2020b). Results showed no significant correlations between any of the four parameters of decoding curves and the level of prediction of behavior (Figure 4B). Therefore, more efficient combinations of features (as measured by higher decoding accuracies) did not correspond to more accurate prediction of behavior.

**Figure 4.**
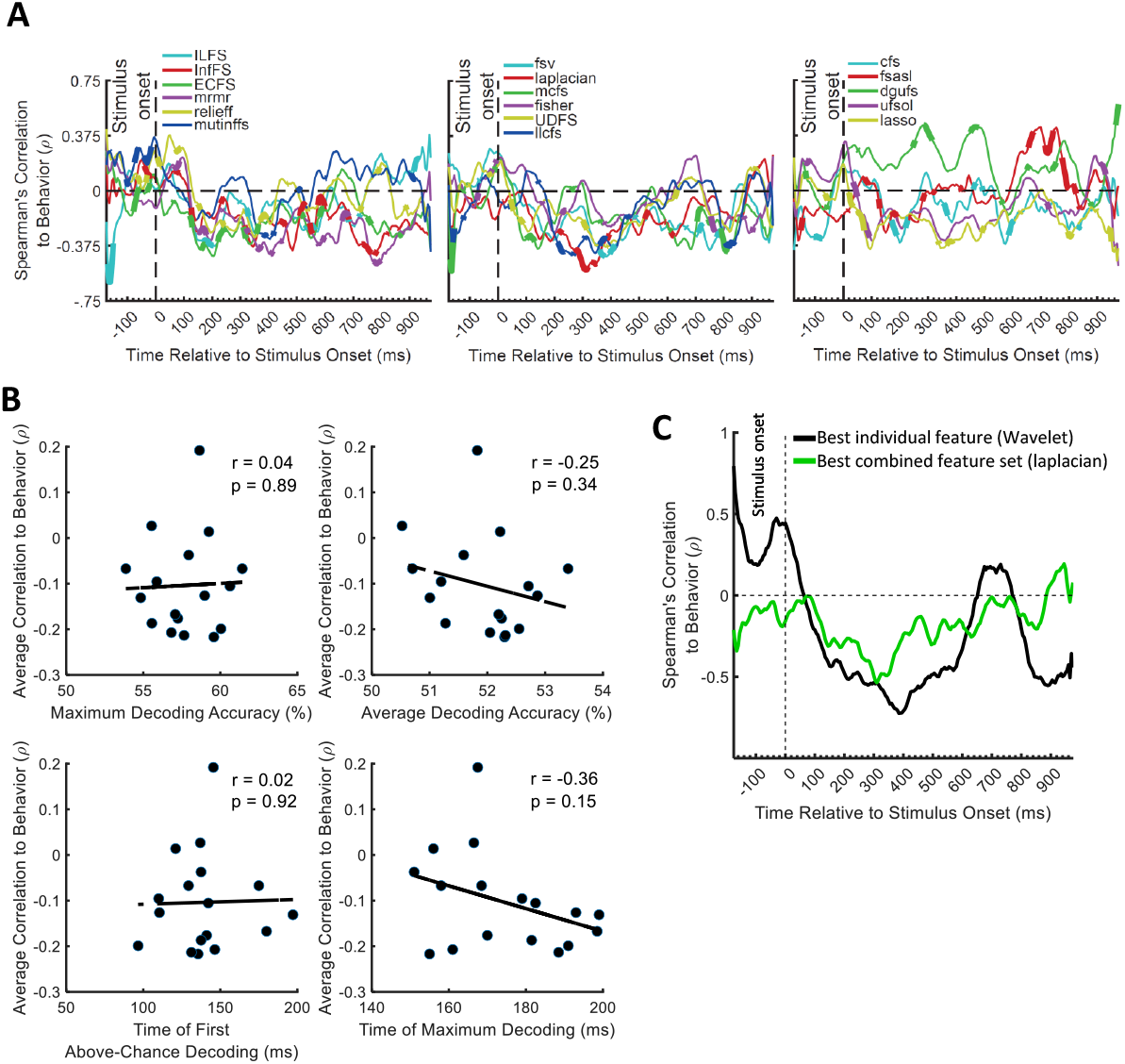
Correlation between the decoding accuracies obtained using 17 feature selection (FS) algorithms and behavioral reaction time of Dataset 2. (A) Top section in each panel shows the (Spearman’s) correlation coefficient obtained from correlating the decoding values and the reaction times for each feature separately. Thickened time points on the curves indicate time points of positively or negatively significant (P<0.05; corrected for multiple comparisons) correlations as evaluated by random permutation of the variables in correlation. (B) Correlation between each of the amplitude and timing parameters of time-resolved decoding (i.e. maximum and average decoding accuracy and time of first and maximum decoding) with the average time-resolved correlations calculated from (A) for the set of N=17 FS algorithms. The slant line shows the best linear fit to the distribution of the correlation data. (C) Correlation between the decoding accuracies obtained from the feature which showed the highest maximum correlation from individual features (Wavelet) and from the combined features (laplacian).

To visually compare the behavioral prediction power of the top-performing individual and combined features we plotted their correlation-to-behavior results on the same figure (Figure 4C). For this we selected Wavelet and laplacian FS, based on them being the single feature and FS algorithm with the largest negative peak. We used this, rather than selecting based on average correlation with behavior because the temporal position of the peak can also provide some temporal indication about the timing of the decision, which if reasonable (e.g. after 200 ms post-stimulus and before the median reaction times of participants: 1146 ms (Karimi-Rouzbahani et al., 2019)), can be more assuring about the existence of true correlation to behavior. The combined features (laplacian) did not provide a negative peak as large as the Wavelet feature, and tended to underperform Wavelet throughout the time course (Figure 4C). Therefore, in contradiction to our hypothesis, the combined features did not provide additional prediction of behavior compared to the individual feature of Wavelet.

## Discussion

Previous studies have used a wide variety of features of neural activations to extract information about visual object categories. However, they have generally used whole-trial analyses, which hide the temporal dynamics of information processing, or time-resolved decoding analyses, or considered the response at each time point separately, ignoring potentially informative temporal features of the time series data. To fill this gap, our previous study extracted and compared a large set of features from EEG in time-resolved analysis (Karimi-Rouzbahani et al., 2021b). However, an outstanding question in the literature was whether the neural code might be best captured by combinations of these features, i.e., if the brain uses a combinatorial encoding protocol to encode different aspects of the sensory input using distinct encoding protocols on the same trial (Gawne et al., 1996; Victor, 2000; Montemurro et al., 2008). Alternatively, previous invasive neural recording studies have suggested a flexible multiscale encoding procedure that allows the generation of all the information within the same platform (Kayser et al., 2009; Panzeri et al., 2010). To address this question we combined a large set of distinct mathematical features (n=26) of the EEG time series data from three datasets, and combined them using a large set of FS algorithms (n=17), each having different criteria for selection. We compared the performance of different FS algorithms using multivariate decoding of category information. Our results showed that, no matter how we combined the informative features, their combined decodable information about object categories, and their power in predicting behavioral performance, was outperformed by the most informative individual feature (i.e. Wavelet), which was sensitive to multi-scale codes from the analysis time window and across electrodes (i.e. spatiotemporal specificity).

The main question of this study was whether the brain recruits and combines a number of different protocols to encode different aspects of cognitive processes involved in object category recognition ranging from sensory information to behavioral response. For example, the brain may use one encoding protocol for the encoding of feed-forward visual information processing, e.g. theta-band power, which would later in the trial be dominated by alpha/beta-band feedback information flow involved in semantic object categorization (Bastos et al., 2015). The brain may also use different encoding protocols to process different aspects of the same stimulus (e.g. contrast or the orientation of visual stimulus; (Gawne et al., 1996)). Alternatively, the brain may implement a single but multiscale protocol (e.g. multiplexing strategy which combines the codes at different time scales (Panzeri et al., 2010)) which allows different aspects of information to be represented within the same encoding protocol. Our results provide support for the latter by showing that spatiotemporally sensitive features, which can detect patterns across multiple scales (e.g. Wavelet coefficients) best capture variance in the EEG responses evoked by different categories of visual objects. Therefore, rather than a combinatorial and switching encoding protocol, the brain may instead encode object category information through a single but multiscale encoding protocol.

This study does not provide the first evidence showing that temporal patterns of activity provide information about different aspects of visual sensory input. The richness of information in the temporal patterns of activity has been previously observed in light encoding (Gollisch and Meister, 2008), co-occurrences of visual edges (Eckhorn et al., 1988), orientations in primary visual cortex (Celebrini et al., 1993) as well as object category information in the temporal cortex (Majima et al., 2014). This study aligns with the recent move towards incorporating within- and across-trial temporal variability in the decoding of information from neural time series such as MEG (Vidaurre et al., 2019), EEG (Majima et al., 2014), invasive electrophysiological (Orbán et al., 2016) and even fMRI (Garrett et al., 2020) data. On the other hand, this current study contrasts with the conventional time-resolved decoding analyses which merely consider amplitude at each time point (Grootswagers et al., 2017), overlooking informative multi-scale temporal codes.

The field of Brain-Computer Interface (BCI) has already achieved great success in decoding visually evoked information from EEG representations in the past two decades, mainly through the use of rigorous supervised learning algorithms (e.g. Voltage Topographies (Tzovara et al., 2011), Independent Component Analysis (Stewart et al., 2014), Common Spatial Patterns (Murphy et al., 2011) and Convolutional Neural Networks (Seeliger et al., 2017)) or by combining multiple features (Chan et al., 2011; Torabi et al., 2017; Wang et al., 2012; Qin et al., 2016). However, the predictive power of a feature about behavior might not be as important for BCI where the goal is to maximize the accuracy of the commands sent to a computer or an actuator. In contrast, one of the most critical questions in cognitive neuroscience to understand whether the neural signatures that we observe are meaningful in bringing about behavior, as opposed to being epiphenomenal to our experimental setup (e.g., Williams et al., 2007; Jacobs et al., 2009; Ritchie et al., 2015; Hebart and Baker; 2018; Woolgar et al., 2019; Karimi-Rouzbahani et al., 2021a; Karimi-Rouzbahani et al., 2021b). To address this point, we evaluated whether our extracted features and their combinations were behaviorally relevant, by correlating our decoding patterns with the behavioral object recognition performance (reaction times in Dataset 2). Moreover, to directly compare the information content of the combined feature sets with the individual features, we equalized the dimensions of the data matrix for the FS algorithm to that obtained for individual features. This avoided artefactualy improving behavioral predictive power with higher dimensionality. Contrary to what we predicted, however, we observed that even the laplacian FS algorithm, which provided the best peak prediction for the behavioral performance, was outperformed by the individual Wavelet feature at most time points. Therefore, the multiscale feature of Wavelet not only provides the most decodable information, but seems to most closely reflect the neural processes involved in generating participant behavior.

One unique property of our decoding pipeline, which we believe led to the enhanced information encoding for the Wavelet feature relative to other individual features (Karimi-Rouzbahani et al., 2021b), is the incorporation of ***spatiotemporal*** codes in decoding in each 50ms analysis window. The neural code can be represented in either time (across the analysis time window), space (across electrodes in EEG) or a combination of both (Panzeri et al., 2010). Specifically, most of the previous studies have evaluated the neural codes in either time, being limited by the nature of their invasive recording modality (Benucci et al., 2009; Houweling and Brecht, 2008), or space by averaging/down-sampling of data within the analysis window. However, our spatiotemporal concatenation of EEG activity across both time and electrodes (i.e. performed at the first PCA stage for individual features and at the third PCA stage for the combined features in Figure 1), allows the neural codes to be detected from both spatially and temporally informative patterns. The 50ms time window chosen here makes a compromise between concatenating and decoding the whole time window in one shot, which loses the temporal resolution, and time-resolved decoding at each time point, which ignores temporal patterns of activity (Karimi-Rouzbahani et al., 2021b).

There are several future directions for this research. First, as the encoding protocols for different cognitive processes might be different from object category processing (Panzeri et al., 2010), the generalization of our results to other domains of cognitive neuroscience needs to be evaluated. Second, previous results (Panzeri et al., 2010) suggest that different aspects of information (e.g. category processing, decision making and motor response) may be encoded using different encoding protocols. Our data did not allow us to tease those aspects apart, which is interesting area for future investigation. Third, following previous suggestions that even different aspects of *visual* information (e.g. color, variations and task) might also be encoded using different encoding protocols (Gawne et al., 1996), the number of selected features might need to be varied from one dataset to another. Ideally, we would only keep the informative features above a certain threshold. Here, we chose an arbitrary threshold of 5 included, but it would be interesting to explore the impact of this parameter in the future.

The large-scale EEG analysis of this study aligns with the recent shift to cross-dataset meta-analyses for different human cognitive abilities such as working memory (Adam et al., 2020) and sustained attention (Langner et al., 2013). Such studies lead to more generalizable conclusions and provide deeper insights into the human cognition. Here, across three very different datasets we showed that, the brain seems to implement a temporally and spatially flexible and multiscale encoding strategy rather than a combinatorial or switching encoding strategy, at least in object category processing.

## Acknowledgements

This research was funded by UK Royal Society’s Newton International Fellowship SUAI/059/G101116 to H.K-R. and MRC intramural funding SUAG/052/G101400 to A.W.

## Authors’ contributions

**Hamid Karimi-Rouzbahani**: Conceptualization, Methodology, Formal analysis, Writing - original draft, Visualization, Data curation, Funding acquisition.

**Alexandra Woolgar**: Writing - review & editing, Funding acquisition.

## Supplementary Materials

**Supplementary Figure 1.**
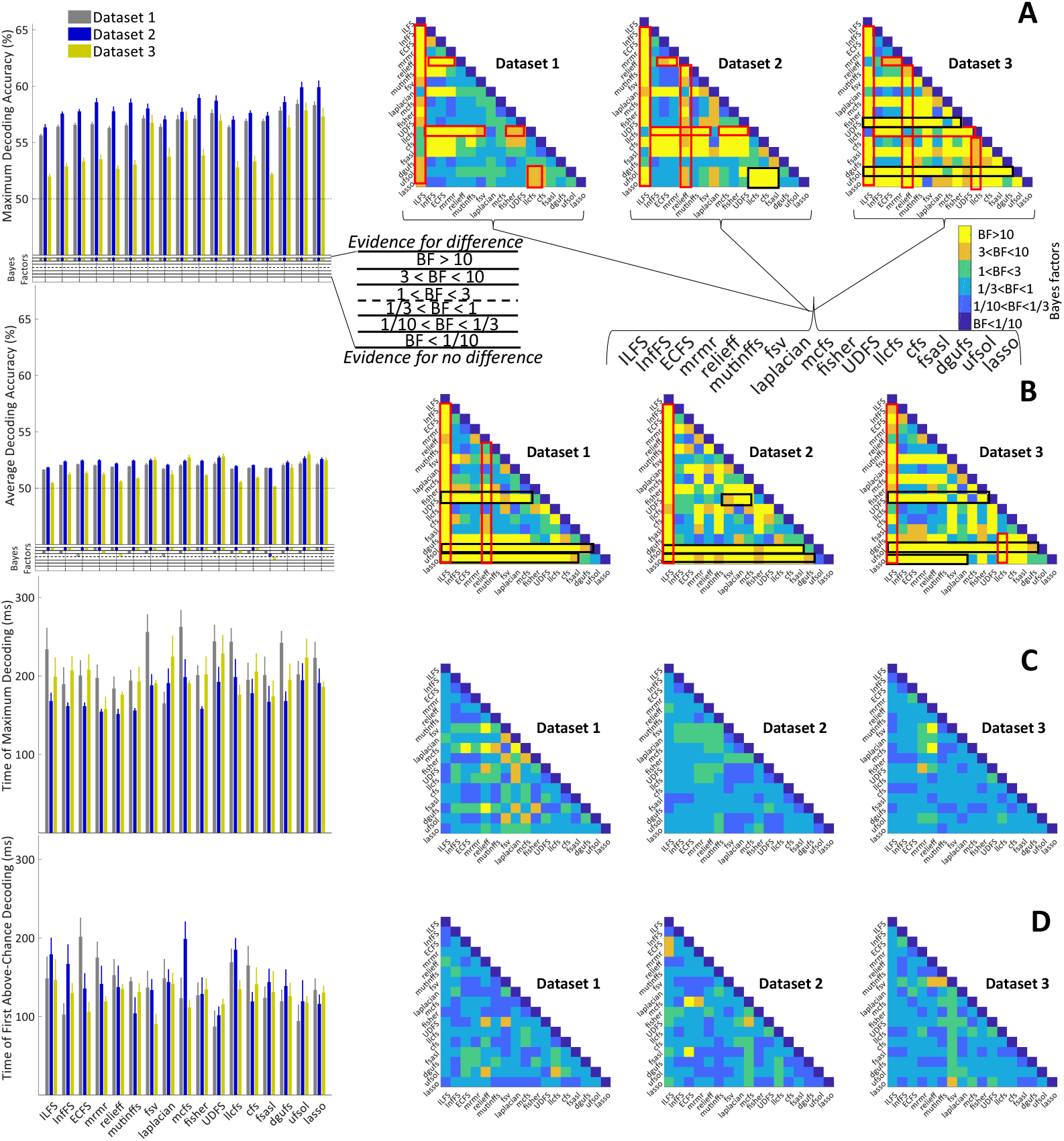
Timing and amplitude parameters extracted from the time-resolved accuracies (Figure 2) of each FS algorithm and each dataset and their Bayesian evidence analyses. (A-D) Left: the maximum and average decoding accuracies, the time of maximum and the first above-chance decoding. Bottom section on A and B show the Bayes factor evidence for the difference of the decoding accuracy compared to chance-level decoding; Right: matrices compare the right parameters obtained from different features. Different levels of evidence for existing difference (moderate 3<BF<10, Orange; strong BF>10, Yellow), no difference (moderate 1/10<BF<1/3, light blue; strong BF<1/10, dark blue) or insufficient evidence (1<BF<3 green; 1/3<BF<1 Cyan) for either hypotheses. Black and red boxes show moderate or strong evidence for higher decoding values for specific features compared other sets of features as explained in the text. The horizontal dashed lines on the left panels of (A) and (B) refer to chance-level decoding. Filled circles in the Bayes Factors show moderate/strong evidence for either hypothesis and empty circles indicate insufficient evidence.

**Supplementary Figure 2.**
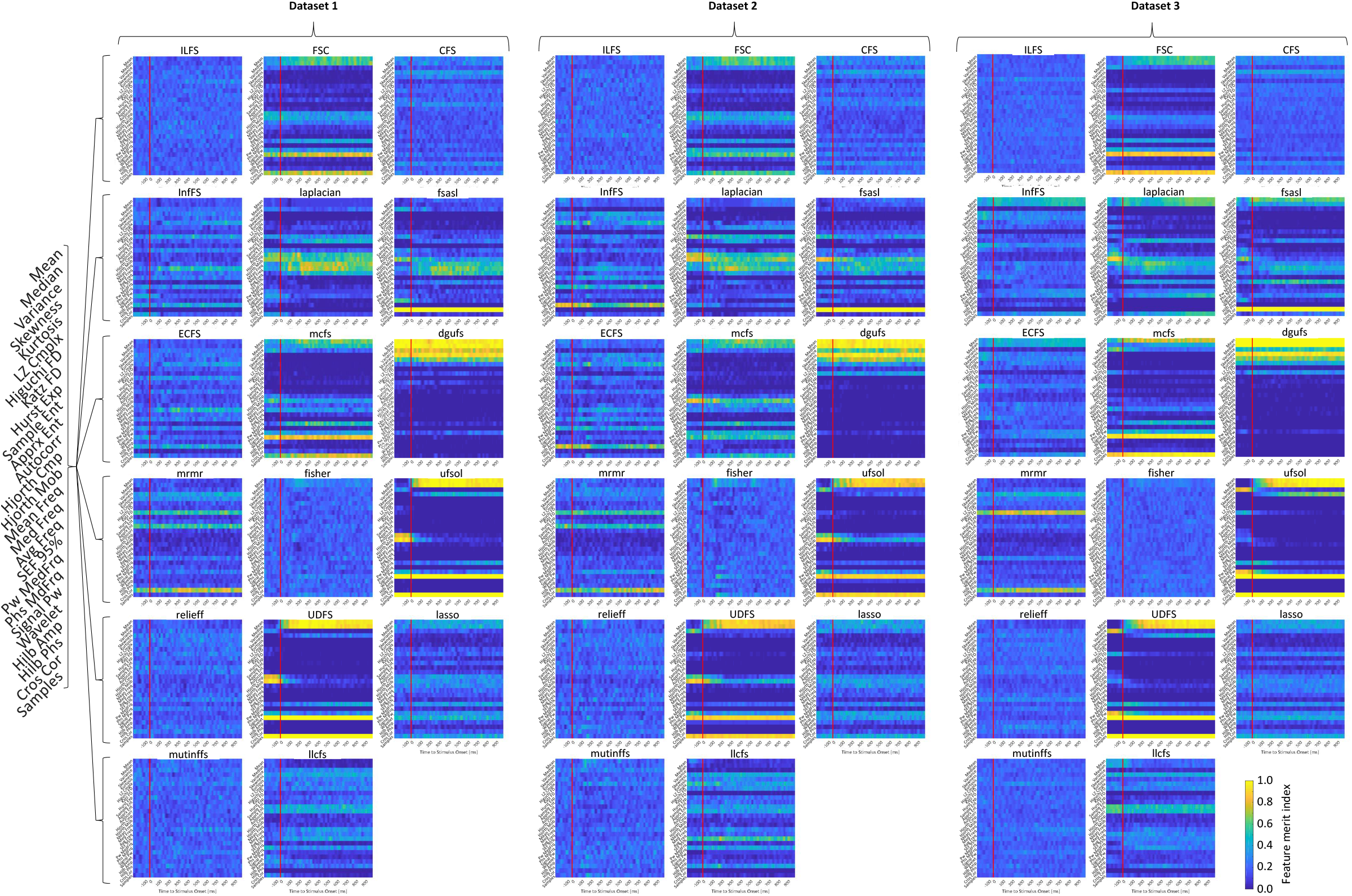
The merit of individual features when combined to maximize the decoding accuracies using 17 different feature selection methods. Warm colors indicate higher merit and cold colors indicate lower merit for the feature at the indicated time point across the trial. Each of the three columns shows the results for one dataset. Merit is the richness of information in the feature about object categories.

## Footnotes

1 https://www.mathworks.com/matlabcentral/fileexchanqe/38211-calc_lz_complexity

2 https://ww2.mathworks.cn/matlabcentral/fileexchange/50290-higuchi-and-katz-fractal-dimension-measures

3 https://www.mathworks.com/matlabcentral/fileexchange/32427-fast-approximate-entropy

4 https://klabhub.github.io/bayesFactor/

